# Dynamics of ion fluxes between neurons, astrocytes and the extracellular space during neurotransmission

**DOI:** 10.1101/305706

**Authors:** N. Rouach, K. Dao Duc, J. Sibille, D. Holcman

## Abstract

Ionic homeostasis in the brain involves redistribution of ionic fluxes in several cell types and compartments, including neurons, astrocytes and the extracellular space. How the major ionic activity-dependent fluxes of potassium and sodium are individually regulated remains difficult to dissociate and to track experimentally. We here review recent progress in modeling the ionic fluxes evoked by neuronal activity based on mass conservation. Excitability of neurons indeed relies on inward sodium and outward potassium fluxes during action potential firing. Recently, we have developed a tri-compartment model based on mass-action kinetics equations that account for potassium dynamics between neurons, astrocytes and the extracellular space. This review describes how such type of model can be used to spatially and temporally predict potassium fluxes during various regimes of neuronal activity. In particular, the model initially showed that it takes several seconds for astrocytes to buffer the majority of the potassium rapidly released by neurons in both basal and high regime of activity. Such model can also probe the selective contribution of ionic channels, and revealed for instance that disruption of the main astroglial potassium K_ir_4.1 channels not only favors the emergence of epileptiform activity, but also dysregulates neuronal excitability specifically during slow rhythmic activities. We here also extend the predictions of the model by assessing the selective contribution of the astroglial and neuronal Na/K ATPase, or volume of the extracellular space on potassium dynamics. We discuss these findings and their implications for neuronal information processing in the healthy and diseased brain.

## Introduction

Modeling fluxes of ions such as potassium (K^+^) and sodium (Na^+^) is a crucial step for understanding the regulation of brain activity at a large scale. It is indeed difficult to dissociate and to track experimentally the flow of ions across various cell types. Astrocytes, the main glial cell type of the central nervous system, form tripartite synapses by enwraping more than half of the synaptic elements with their peripheral processes in the hippocampal CA1 region (Bushong et al., 2004; Perea et al., 2009). Thanks to their proximity to synapses, astrocytes can detect and process neuronal activity via multiple pathways, including calcium signaling (Verkhratsky et al., 2012) and ionic currents mediated by transporters or ionic channels, activated by GABA (gamma-aminobutyric acid), glutamate (Bergles and Jahr, 1997; Diamond et al., 1998; Luscher et al., 1998; Goubard et al., 2011) or K^+^ (Orkand et al., 1966; Karwoski et al., 1989; Meeks and Mennerick, 2007). Astroglial cells not only process, but also modulate neuronal activity by various mechanisms involving intracellular signaling or extracellular homeostasis, in particular the regulation of the extracellular space volume or the levels of GABA, glutamate or K^+^ (Nedergaard and Verkhratsky, 2012). Remarkably the first investigations focusing on astrocytic-evoked signals following neurotransmission was reported in the frog optic nerve as a membrane potential depolarization due to K^+^ entry across the astrocytic membrane (Orkand et al., 1966). This was suggested to contribute to K^+^ spatial buffering, a process consisting in astroglial local uptake of excess extracellular K^+^ ([K^+^]_o_) following neuronal activity, redistribution in the astroglial networks mediated by gap junction channels and release at distal sites of low [K^+^]_o_ (Kuffler et al., 1966). Several astroglial K^+^ channels and transporters participate to the extracellular clearance of K^+^, including inward rectifier 4.1 (K_ir_4.1) two-pore K^+^ channels (K_2P_) and Na/K ATPases (Walz, 2000; Butt and Kalsi, 2006). Besides non selective blockers of K_ir_ channels, recently the use of conditional glial K_ir_4.1 knockout mice (GFAP-Cre-K_ir_4.1fl/fl mice, K_ir_4.1^−/−^) has suggested an important role for K_ir_4.1 channels in the astrocytic control of [K^+^]_o_ (Djukic et al., 2007; Chever et al., 2010; Haj-Yasein et al., 2011; Bay and Butt, 2012). However K_ir_4.1 deletion throughout life in glial cells, including astrocytes but also oligodendrocytes (Djukic et al., 2007), leads to major brain defects and possibly compensations that likely obscure the primary role of K_ir_4.1 channels in basal neurophysiology. Astrocytes from K_ir_4.1^−/−^ mice are indeed strongly depolarized (Djukic et al., 2007; Chever et al., 2010; Haj-Yasein et al., 2011) and these mice display not only ataxia, seizures, white matter vacuolization or growth retardation, but also die prematurely about three weeks after birth. It is thus challenging to assess experimentally the selective contribution of astrocytic K_ir_4.1 channels to the moment to moment regulation of [K^+^]_o_ and neuronal activity. An alternative approach to assess such contribution consists in computational modeling. However, most modeling studies have focused on astroglial dysregulation of [K^+^]_o_ in pathological conditions to investigate its role in the generation of epileptiform activity. Alterations in astroglial regulation [K^+^]_o_ homeostasis indeed contribute to seizures by controlling both their initiation and maintenance (Cressman et al., 2009; David et al., 2009; Ullah et al., 2009). We here review several of these models and in particular a tri-compartment model we recently developed (Sibille et al., 2015), which temporally predicts K^+^ dynamics between neurons, astrocytes and the extracellular space during various regimes of neuronal activity. The latter is based on synaptic dynamics, Hodgkin-Hukley model of excitability and intercellular flux balance, and has already provided several key results regarding the contribution in physiological conditions of K_ir_4.1 astroglial channels on [K^+^]_o_ and neurotransmission. We here also extend these results by assessing the selective contribution of the astroglial and neuronal Na/K ATPases and of the volume of the extracellular space on K^+^ dynamics.

## Recent tripartite models

A variety of computational models have investigated extracellular K^+^ regulation of neurotransmission, including glial uptake mechanisms (Kager et al., 2000; Dronne et al., 2006; Dronne et al., 2007; Cressman et al., 2009; David et al., 2009; Florence et al., 2009; Ullah et al., 2009; Oyehaug et al., 2012). To investigate mechanisms of seizure discharges and spreading depression, an initial tri-compartment model which includes the astrocytes, neurons and extracellular space was performed (Kager et al., 2000). In this model, the astrocyte membrane potential was not considered, and the accumulation of K^+^ in the interstitial volume was set by a first-order buffering scheme that simulates glial K^+^ uptake. This model showed that after evoked firing, it takes ~17 s for the membrane potential of neurons to return to resting values, and this occurs solely via activation of Na/K ATPases. This model also predicts that high [K^+^]_o_ play an important role in the initiation and maintenance of epileptiform activity.

Another tri-compartment model, simplified as a one-dimensional two-layer network, has been performed to assess how networks of neurons switch to an activity persistent state, and the stability of the persistent state to perturbations (Cressman et al., 2009). In this model, Na^+^ and K^+^ alter neuronal excitability, frequency of seizures, and stability of activity persistent states. This model also provides the quantitative contribution of intrinsic neuronal currents, Na/K ATPases, glia, and diffusion of extracellular Na^+^ and K^+^ to slow and large-amplitude oscillations in extracellular and neuronal Na^+^ and K^+^ levels. The levels of [K^+^]_o_ estimated during epileptiform activity were comparable to the ones observed experimentally (McBain, 1994; Ziburkus et al., 2006). Although this model does not account for astrocytic K_ir_4.1 channels, it already shows that a local persistent network activity requires both, balanced excitation and inhibition, and glial regulation of [K^+^]_o_ (Ullah et al., 2009).

Finally, a model taking into account the extracellular space and astroglial compartments has provided a quantitative description of the contribution of several astrocytic ionic channels and transporters (NKCC1, Na/K ATPase, NBC, K^+^, Na^+^ and AQ channels) in the modulation of neuronal excitability (Oyehaug et al., 2012).

## Modeling activity-dependent potassium dynamics between neurons, astrocytes and the extracellular compartment

Recently we developed a model of K^+^ ions dynamics during neuronal activity at a biophysical level that includes neurons, astrocytes and the extracellular space (Fig. 1). As previously described (Sibille et al., 2015), neuronal cells are approximated by a single compartment (Kager et al., 2000; Dronne et al., 2006; Cressman et al., 2009) with a membrane equipped with conductance-based Na^+^ and K^+^ voltage-gated channels. The neuronal membrane potential is coupled to the intracellular and extracellular Na^+^ and K^+^ concentration by the currents. In contrast with previous approaches (Kager et al., 2000; Cressman et al., 2009; Ullah et al., 2009; Oyehaug et al., 2012), we include explicitly in the model the astrocytic membrane potential dynamics, which we also experimentally measured under different stimulation patterns. This dynamics is directly coupled with the K^+^ flux associated with K_ir_4.1 activity.

**Figure 1.**
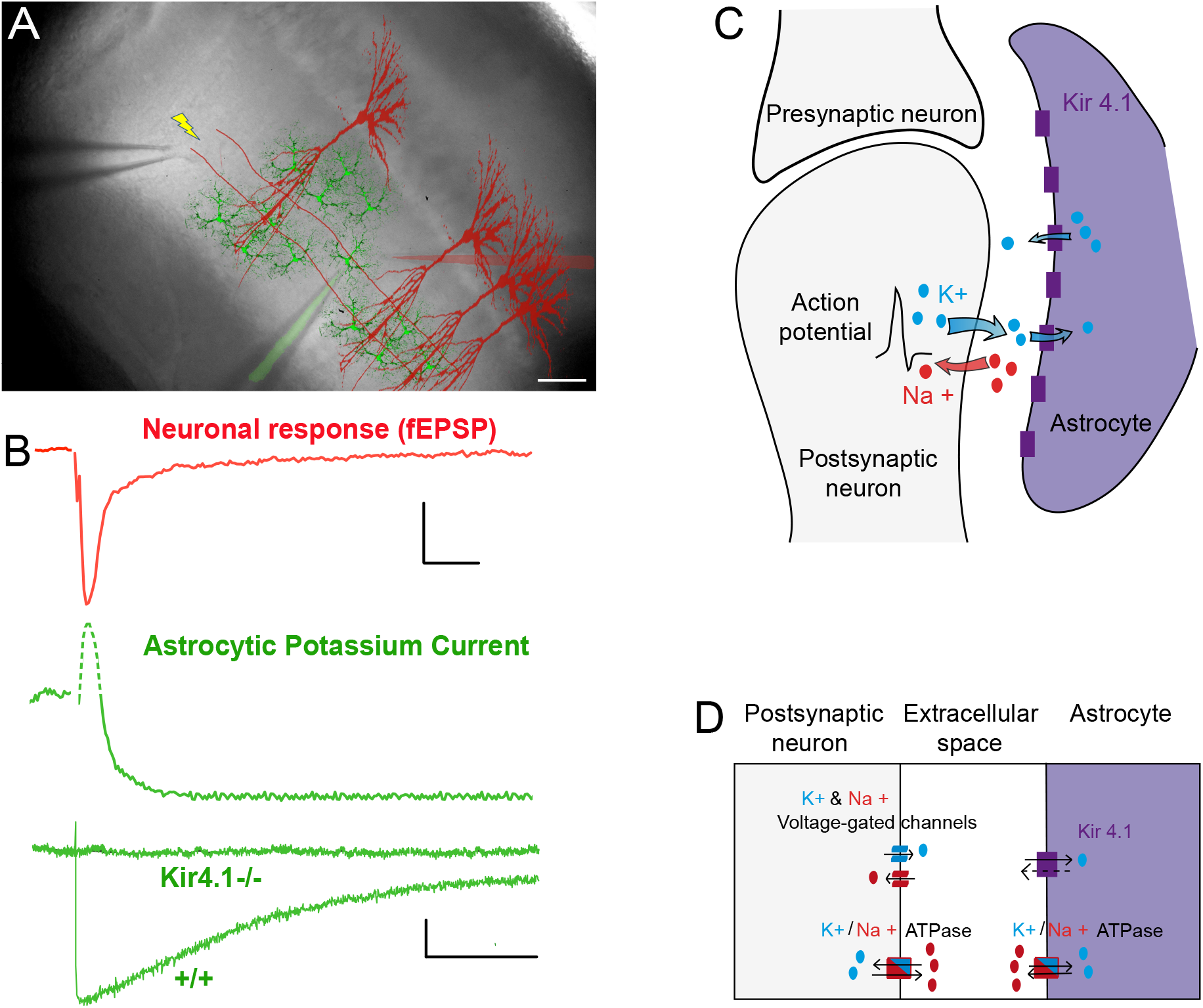
Synaptically-evoked potassium current in astrocytes and tri-compartment model of the potassium cycle between neurons, extracellular space and astrocytes. *A)* Sample scheme of the hippocampal CA1 region illustrating the arrangement of the stimulating electrode (left) to activate (yellow lightning) the Schaffer collaterals (SC, red lines), the patch pipette electrode (filled in green), to record currents from astrocytes (green), and the neuronal recording electrode (extracellular pipette, filled in red), to record fEPSP generated by CA1 pyramidal cells (red). Scale bar, 50 μm. *B)* Sample traces (2 upper traces) of synchronous recording of neuronal responses (field excitatory postsynaptic potential (fEPSP, red) and pharmacologically isolated astroglial whole-cell K^+^ currents - obtained by subtracting the kynurenic acid (5 mM) insensitive component from the total current recorded with 100 μM picrotoxin - induced by SC stimulation in the hippocampal CA1 region. Scale bar, 0.1 mV for fEPSP, 15 pA for astroglial current, 25 ms. Note that the initial fast outward astroglial current component (dash green line) reflects the fEPSP, while the astroglial K^+^ current is the subsequent slow inward component (solid green line). Sample traces (lower superimposed traces, green) of the pharmacologically isolated astroglial K^+^ current in wild type and K_ir_4.1 glial conditional knockout mice (Kir4.1^−/−^). This current is fully abolished in K_ir_4.1^−/−^ mice, showing the sole contribution of K_ir_4.1 channels the isolated astroglial K^+^ current. Scale bar, 15 pA, 2 s. *C, D*) Tri-compartment model accounting for the K^+^ cycle between the neuron, the extracellular space and the astrocyte. *C*) Schematic diagram describing the tri-compartment model where evoked neuronal activity releases K^+^ extracellularly, which is subsequently taken up by neighboring astrocytes. *D*) Simplification of the tri-compartment model to ionic fluxes exchanges between a postsynaptic neuron, a perisynaptic astroglial process and the extracellular space. The model includes channels and pumps carrying K^+^ and Na^+^ ions.

The ion concentrations depend on the activity of neuronal and astroglial Na/K ATPases, which maintain the resting [K^+^]_i_ by balancing the K^+^ and Na^+^ fluxes. Dynamics in the astrocytes was modeled by a single compartment equipped with a membrane including conductance-based astrocyte containing K_ir_4.1 channels. K_ir_4.1 channels are inward rectifier K^+^ channels strongly expressed in hippocampal astrocytes and that primarily generate the long-lasting synaptically-evoked astroglial K^+^ currents, as shown experimentally using dual electrophysiological recordings of neuronal and astrocytic responses in wild type and K_ir_4.1^−/−^ mice (Dallerac et al., 2013) (Fig. 1A,B).

We now summarize this model where neurons and astrocytes are separated by an extracellular compartment. The model is based on balancing ionic fluxes between the three compartments (Fig. 1C, D).

The dynamics of the synaptic response of a postsynaptic neuron ismodeled by a current I*app*, used as the initial input of a classical Hodgkin-Huxley model, accounting for the neuronal membrane potential dynamics (entry of Na^+^ and exit of K^+^). Released extracellular K^+^ are then taken up by astrocytes via K_ir_4.1 channels and Na/K ATPases (Fig. 1D).

K_ir_4.1 channels are involved in K^+^ uptake (Djukic et al., 2007). To capture this effect, the biophysical properties of this channel, i.e. the current-voltage relationship (IV curve) of K^+^ ions through K_ir_4.1 channels (Fig. 2A), can be used according to equation 22. Furthermore, our model predicts how this IV curve changes for various values of [K^+^]_o_ (Fig. 2B). Interestingly, K^+^ fluxes through K_ir_4.1 channels vanish around astrocytic resting membrane potential (~ −80 mV) and are outward during astrocytic depolarization for a fixed [K^+^]_o_ (2.5 mM, Fig. 2A). However, they become inward when [K^+^]_o_ increases (5−10 mM, Fig. 2B).

**Figure 2.**
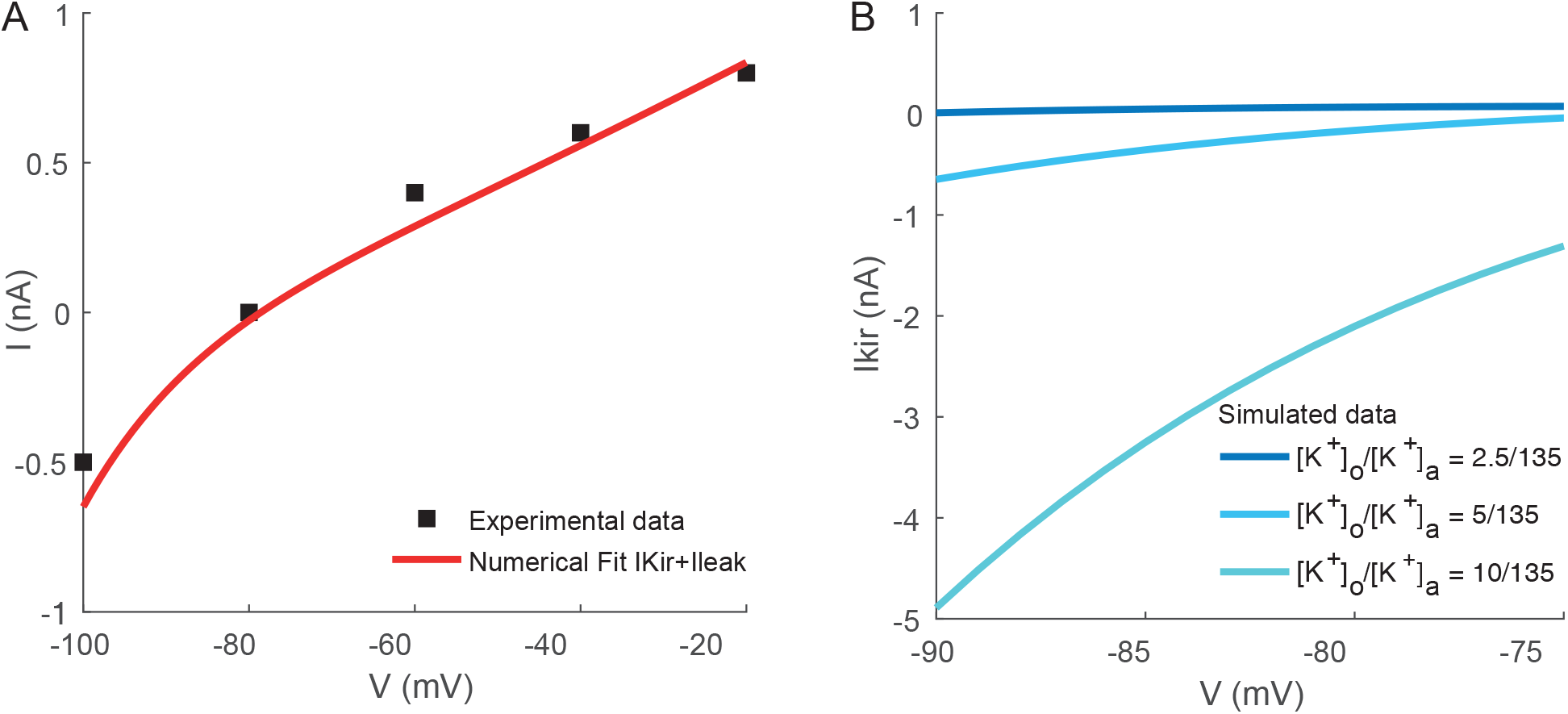
Fitting and simulations of the biophysical properties of Kir 4.1 channels. *A)* Illustration of current-voltage (IV) relationship of K_ir_4.1 channels recorded experimentally (from Fig. 4 Ransom 95, black) and our corresponding numerical fit for [K^+^]_o_ = 3 mM (red) and [K^+^]_a_ = 135 mM, using equation (22) *B)* K^+^ dependence of the different simulated IV curves of *I_Kir_* ((Eq. 22) for [K^+^]_o_ = 2.5, 5 or 10 mM.

## Modeling equations for the potassium fluxes between neuron/astrocyte/extracellular space

The biophysical model to describe K^+^ dynamics during neuronal activity and specifically the role of astroglial K_ir_4.1 channels consists of a set of equations that describe the membrane potential and concentration changes following stimulations of the Schaffer collaterals (SC), presynaptic release of glutamate and activation of postsynaptic neurons. This step is modeled by the classical facilitation/depression model (Tsodyks and Markram, 1997). The resulting postsynaptic activity triggers release of ions in the extracellular space and a change in the astrocytic membrane potential via ion uptake.

### Facilitation-Depression model

To account for SC stimulation, which induces a postsynaptic response in the hippocampal CA1 *stratum radiatum* region, we used a facilitation-depression model (Tsodyks and Markram, 1997; Markram et al., 1998; Tsodyks et al., 1998).

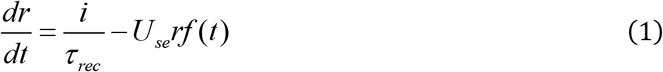

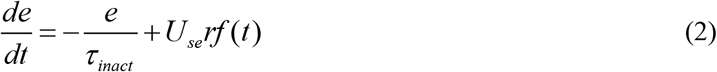

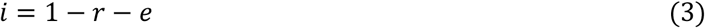

where *f* is defined as the input function. For a single stimulation generated at time *t_stim_, f*(*t*) = *δ*(*t* – *t_stim_*).

A stimulation instantaneously activates a fraction *U_se_* of synaptic resources *r*, which then inactivates with a time constant *τ_inac_* and recovers with a time constant *τ_rec_*. In the simulations, at time *t* = *t_stim_, r* and *e* respectively decreases and increases by the value *U_se_r*. The synaptic current *I_app_* is proportional to the fraction of synaptic resources in the effective state *e* and is given by *I_app_* = *A_se_e* (the parameter *A_se_* is defined in table 1). We used the following definitions for the input function *f*:

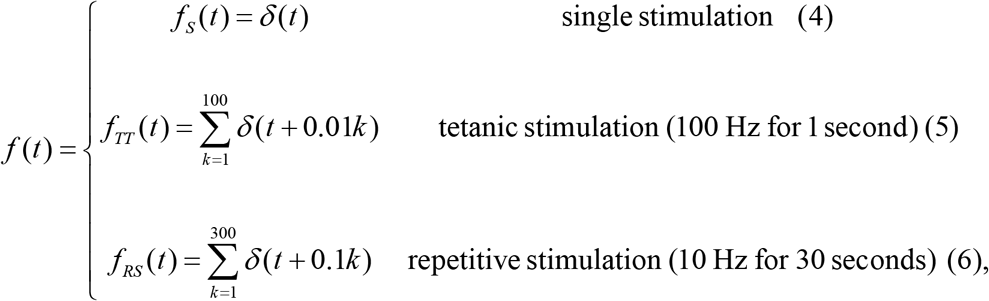

**Table 1.** Parameters of the model.

|  | Parameters | Value |
| --- | --- | --- |
| $\tau_{rec}$ | Recovery time constant | 300 ms (WT) / 500 ms (fitted $K_{ir}4.1^{-/-}$) (Tsodyks and Markram, 1997) |
| $\tau_{inac}$ | Inactivation time constant | 200 ms (WT) / 160 ms (fitted $K_{ir}4.1^{-/-}$) (Tsodyks and Markram, 1997) |
| $A_{se}$ | Absolute synaptic strength | 7 (WT) / 10 (fitted $K_{ir}4.1^{-/-}$) (Tsodyks and Markram, 1997) |
| $U_{se}$ | Utilization of synaptic efficacy | 0.8 (WT) / 0.8 ($K_{ir}4.1^{-/-}$) |
| $g_{Na}$ | Neuronal sodium channel conductance | 15 nS |
| $g_K$ | Neuronal potassium channel conductance | 4 nS |
| $V_{rest}$ | Neuronal resting membrane potential | -60 mV (Hodgkin et al., 1952) |
| $Na$ | Avogadro Number | $6.02 \times 10^{23}$ |
| $q_e$ | Net charge of single monovalent ion | $1.62 \times 10^{-19}$ C |
| $F$ | Faraday Constant | $9.64 \times 10^{-4}$ C.mol<sup>-1</sup> |
| $R$ | Gaz constant | $8.314$ J.mol<sup>-1</sup>.K<sup>-1</sup> |
| $T$ | Temperature | 308 K |
| $g_{lN}$ | Neuronal leak conductance | 0.07 nS |
| $V_{lN}$ | Neuronal leak potential | -1.2793 mV |
| $C_N$ | Neuronal membrane capacitance | 136 pF (Pannasch et al., 2014) |
| $G_{K_{ir}}$ | Astrocytic single channel $K_{ir}$ conductance | 3.64 nS |
| $V_{A1}$ | $K_{ir}$ current potential constant 1 | + 14.83 mV extracted from (Ransom and Sontheimer, 1995) |
| $V_{A2}$ | $K_{ir}$ current potential constant 2 | -105.82 mV extracted from (Ransom and Sontheimer, 1995) |
| $V_{A3}$ | $K_{ir}$ current potential constant 3 | 19.23 mV extracted from (Ransom and Sontheimer, 1995) |
| $C_A$ | Astrocytic capacitance | 15 pF (Chever et al., 2014) |
| $V_{lA}$ | Astrocytic leaking potential | -74 mV |
| $g_{lA}$ | Astrocytic leak conductance | 0.015 nS |
| $i_{maxA}$ | Astrocytic Na/K pump rate | 0.3 mM.ms<sup>-1</sup> |
| $i_{maxN}$ | Neuronal Na/K pump rate | 0.9 μM.ms<sup>-1</sup> |
| $\frac{Vol_o}{Vol_N}$ | Extracellular space volume/neuronal volume | 0.5 (Chen and Nicholson, 2000) |
| $\frac{Vol_o}{Vol_A}$ | Extracellular space volume/astrocytic volume | 0.5 (Chen and Nicholson, 2000) |
| $i_{NaIN}$ | Neuronal sodium leak rate | $-1.35.10^{-4}$ mM.ms<sup>-1</sup> |
| $i_{NaIA}$ | Astrocytic sodium leak rate | $-1.6.10^{-3}$ mM.ms<sup>-1</sup> |

### Modeling Neuronal Activity

The dynamics of the neuronal membrane potential, *V_N_*, follows the Hodgkin Huxley (HH) equations (Hodgkin and Huxley, 1952).

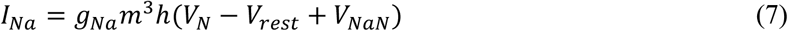

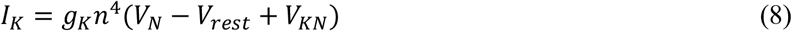

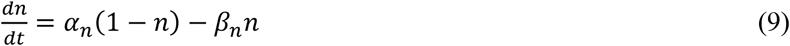

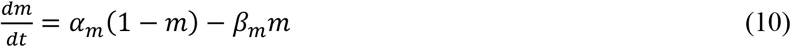

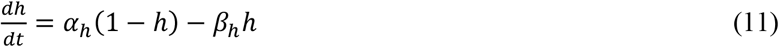

with rate equations

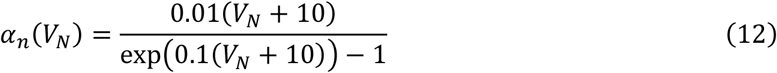

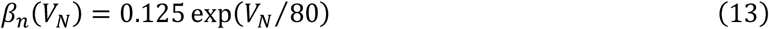

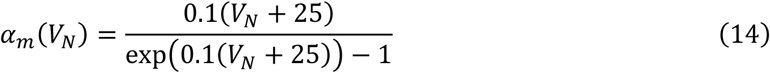

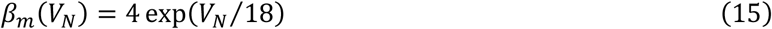

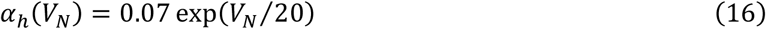

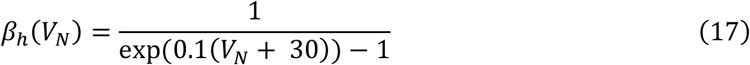

*V_rest_* defines the resting membrane potential and *V_KN_* and *V_NaN_* are respectively the K^+^ and Na^+^ equilibrium potentials and are given by the Nernst equations

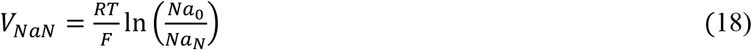

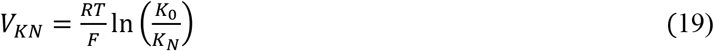

where *Na*_0_ and *Na_N_* are respectively the extracellular and neuronal Na^+^ concentrations, and *K*_0_ and *K_N_* are respectively the extracellular and neuronal K^+^ concentrations that may vary. The leak current is

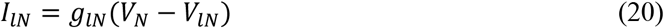

which stabilizes the membrane potential at its resting value. Finally, the neuronal membrane potential satisfies the equation

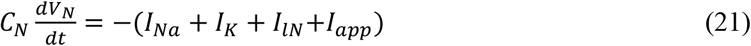

where *I_app_* is the synaptic current derived from equation 1.

### Modeling astrocytic potassium uptake by Kir4.1 channels

To account for the K^+^ dynamics in astrocytes, the K_ir_4.1 channel is considered by using its biophysical properties (Siegenbeek van Heukelom, 1994) and I-V curve (Ransom and Sontheimer, 1995). The total astroglial current *I_Kir_* depends on the membrane potential, the extracellular (*K*_0_) and the astrocytic (*K_A_*) K^+^ concentrations, and is approximated by

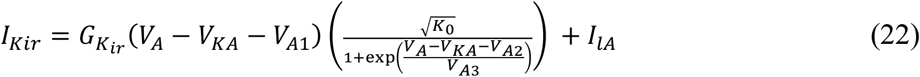

where V_KA_ is the Nernst astrocyte K^+^ potential, V_A_, the astrocyte membrane potential, K_0_ is the extracellular K^+^ concentration and V_A1_ (an equilibrium parameter, which sets K_ir_ current to 0 at −80 mV), V_A2_ and V_A3_ are constant parameters calibrated by the I-V curve (Fig. 2, (Ransom and Sontheimer, 1995)), as detailed below. The leak current I_lA_ = g_lA_(V_A_ – V_lA_) is added to stabilize the astrocyte membrane potential at − 80 mV and contributed to the fit (Fig. 2A).The first term of equation 22 describes the dependence of I_Kir_ to the square root of K_0_ (Hagiwara and Takahashi, 1974; Sakmann and Trube, 1984; Carmeliet et al., 1987; Mazzanti and DeFelice, 1988; Ransom and Sontheimer, 1995) and to the steady state open/close partition function of K_ir_4.1 channels according to the Boltzmann distribution (Siegenbeek van Heukelom, 1994), which includes dynamic variations of K^+^ Nernst potential during neuronal activity.

The astrocyte membrane potential *V_A_* satisfies the equation

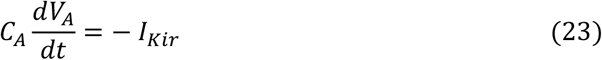

The K_ir_4.1 channel I-V curve (equation 22) was used to fit *I_Kir_* to experimental recordings for the K_ir_4.1 channel (3 mM [K^+^]_o_ (Fig. 2A from Fig. 4 in (Ransom and Sontheimer, 1995; Ransom et al., 1996)), leading to the values *V*_*A*1_ = +14.83 *mV,V*_*A*2_ = –105.82 *mV* and *V*_*A*3_ = 19.23 *mV* (Table 1). It is to be noted that *G_Kir_* = 3.64 *nS*, and g_lA_ = 0.015*nS* (table 1).

### Na/K ATPase ionic flux for astrocytes and neurons

The K^+^ resting concentrations in neurons and astrocytes are maintained by Na/K pumps that balance the outward K^+^ and inward Na^+^ fluxes. The associated pump currents *i_pump,k_* (index *k* = *N* for the neuron, *k* = *A* for the astrocyte) depend on the extracellular K^+^ *K*_0_ and intracellular Na^+^ concentrations (*Na_N_* for the neuron and *Na_A_* for the astrocyte) and follow the same equation as (Reichenbach et al., 1992),

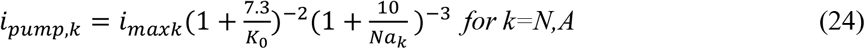

where *i_maxk_* is a constant (Table 1).

### Balance of ionic fluxes

The electrogenic neuronal and astrocytic channel currents can be converted into ionic fluxes (Cressman et al., 2009): a current *I* across a membrane induces a flow of charge *i* equals to *δQ* = *I* per unit of time. The corresponding change in extracellular concentration is given by *I*/(*qN_A_Vol*_0_), where *q* = 1.6 * 10^−19^*C* is the charge of an electron, *N_A_* the Avogadro number and *Vol_N_, Vol_A_* and *Vol*_0_ are the neuronal, astrocytic and extracellular volume respectively. To model the ionic concentration dynamics, the currents *I_Na_, I_K_* and *I_Kir_* are converted into the corresponding ionic fluxes *i_Na_, i_K_* and *i_Kir_*.

### Potassium fluxes

The mass conservation law for the extracellular *K*_0_, the neuronal *K_N_* and the astrocytic *K_A_* K^+^ concentrations are used to determine the system of equations for the K^+^ fluxes: the extracellular K^+^ *K*_0_ increases with the neuronal current *I_K_* (see equation 8), which is here converted to *i_K_* (ion flux), but it is also uptaken back into neurons with a flux 2 *i_pumpN_* (the factor 2 is described in (Cox and Helman, 1986) and into astrocytes as the sum of the two fluxes 2 *i_pumpA_* plus *i_Kir_*. Using the similar equations for the neuronal and astrocytic K^+^ to balance the various fluxes, we have

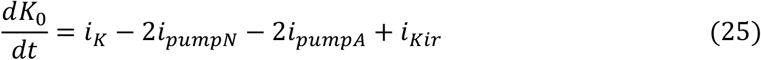

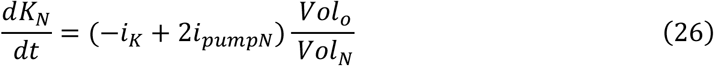

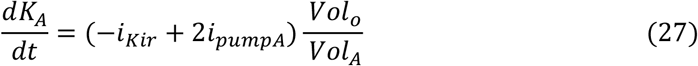

### Sodium fluxes

The equations for the Na^+^ fluxes are derived by the balance between the neuronal, astrocytic and extracellular concentrations. However, the main differences are that the pump exchanges 2 K^+^ for 3 Na^+^ ions, leading to the coefficient 3 in front of the pump term. In addition, to stabilize the Na^+^ concentrations, two constant leak terms *i_NalA_* and *i_NalN_* (values given in table 1) are added (Kager et al., 2000),

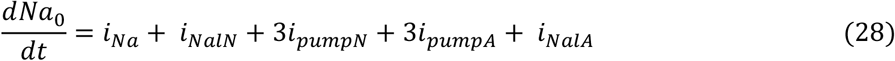

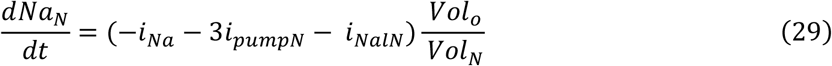

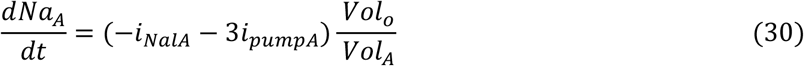

#### Modeling facilitation/depression

To account for the synaptic properties of CA1 pyramidal neurons following single, tetanic and repetitive stimulations, a synaptic current using the depression-facilitation model (equation 1) is generated where *I_app_* depends on the input functions *f_s_*(*t*) (equation 4), *f_TT_*(*t*) (equation 5) and *f_RS_*(*t*) (equation 6), respectively. The parameters for the K_ir_4.1 inhibition condition in the model are extracted from our experimental results on K_ir_4.1 glial conditional knockout mice (Sibille et al., 2014) and are given by *τ_rec_* = 500 *ms, τ_inact_* = 160 *ms*.

#### Simulation of neuronal firing at various frequencies

We imposed an initial input at various frequencies (0.1, 1, 3, 5, 10, 50 Hz). Each input is generated by a sub-firing square current lasting 5 ms (I*app*). In addition, we added a Brownian noise of amplitude *σ* = 0.68 *pA*^2^.*ms*^−1^ to represent neuronal membrane potential fluctuation (equation 21), the amplitude (1 mV) of which was chosen to induce a probabilistic firing of 0.2, matching the CA1 pyramidal cells synaptic release probability p = 0.2 (probability to induce a postsynaptic response in equation 1) (Bolshakov and Siegelbaum, 1995). Using the tri-compartment model, we simulated at various frequencies a quantity that we called the observed firing probability defined empirically at time *t* as the time dependent ratio of the number of spikes observed at time *t* to the total number of simulations.

## Activity-dependent control of extracellular K^+^ clearance, membrane potential dynamics and neuronal excitability by astroglial K_ir_4.1 channels

To investigate the acute role of astroglial cells in extracellular K^+^ homeostasis controlling neuronal activity, we previously combined electrophysiological recordings with the tri-compartment model described above, which accounts for K^+^ dynamics between neurons, astrocytes and the extracellular space (Sibille et al., 2015). It remains difficult to obtain analytical solutions from the 30 equations of the model. However, it can be used to explore the K^+^ neuroglial interactions for various regimes of activity. After generating a synaptic current (I*app*) using the depression-facilitation model (equation 1) (f(t) = δ(t)) combined to the Hodgkin-Huxley model, this synaptic current generates action potential discharges, resulting in firing during the stimulations (1s 100 Hz for tetanic stimulation, 30 s,10 Hz for repetitive stimulation). This induces a transient increase of ~ 0.9 mM [K^+^]_o_ which peak at 3.4 mM within 300 milliseconds of a single stimulation, while [K^+^]_o_ peak at ~ 6.9 mM after 17.5 seconds of repetitive stimulation (Fig. 3A).

**Figure 3.**
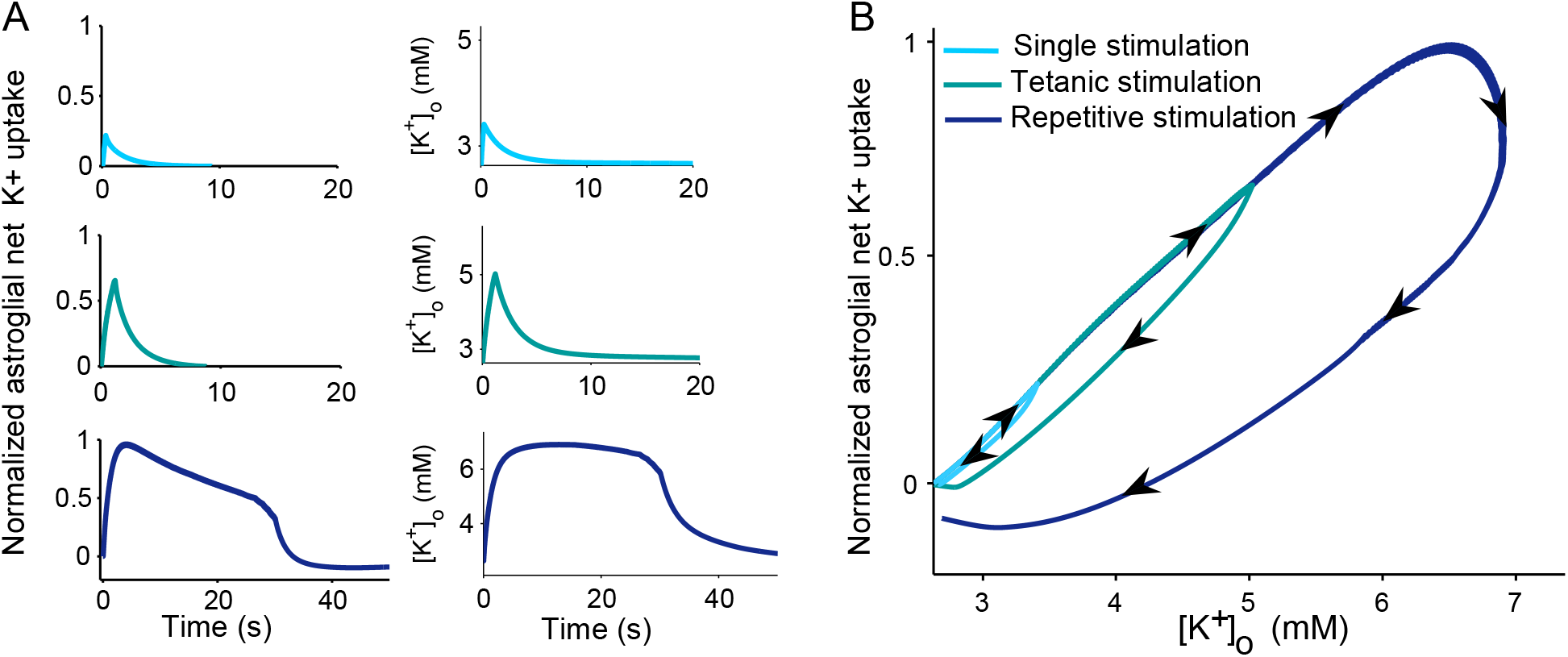
Extracellular K^+^ levels set the strength of astroglial potassium uptake. *A*) Activity-dependent astrocytic net K^+^ uptake (equation 27) and changes in [K^+^]_o_ induced by single (cyan), tetanic (100 Hz, 1s, green) and repetitive (10 Hz, 30 s, dark blue) stimulations are plotted as a function of time. *B)* Phase diagram showing astrocytic K^+^ uptake’s dynamics as a function of activity-dependent changes in [K^+^]_o_ for the three different regimes of evoked activity. Astrocytic K^+^ net uptake is normalized to its maximum value obtained during repetitive stimulation.

We found that activity-dependent transient rise in [K^+^]_o_ set the strength of astroglial net K^+^ uptake, as shown in figure 3, which illustrates the dynamic net K^+^ uptake in astrocytes according to the corresponding variations in [K^+^]_o_ during different regimes of activity, i.e. single, repetitive and tetanic afferent stimulations (Sibille et al., 2015). These data indicate that the astroglial net K^+^ uptake follows the activity-dependent [K^+^]_o_ transient amplitudes (Fig. 3A,), i.e. increases proportionally to the variations in [K]_o_ evoked by the different regimes (Fig. 3B). We also found that astroglial cells do buffer within 6 to 9 seconds more than 80 % of the K^+^ released by neurons in response to basal, repetitive and tetanic afferent stimulations (Sibille et al., 2015). However, the astroglial K^+^ uptake then decreases differentially over time, in a non-linear manner, according to the regime of stimulation (Fig. 3B). Indeed, the time needed for astroglial cells to buffer the released K^+^ is not proportional to [K^+^]o rises, as illustrated by the phase diagram which represents astrocytic K^+^ clearance as a dynamic function of activity-dependent changes in [K^+^]o induced by the different regimes of activity (Fig. 3B). The time it takes for astrocytes to buffer K^+^ is actually proportional to the square root of [K^+^]_o_ (equation 22).

Using such approach, we found that astrocytic K_ir_4.1 channels not only play a major role in extracellular K^+^ clearance, but also in the dynamics of astrocytic and neuronal membrane potentials, in particular during trains of stimulation, and strongly modulate excitability of neurons for theta rhythmic activity (Sibille et al., 2015). We indeed demonstrated that astrocytic K_ir_4.1 channels, besides being crucially involved in the clearance of extracellular K^+^, are sufficient to mediate the synaptically-evoked slow membrane depolarization in hippocampal astrocytes during low and high regimes of neuronal activity. Astrocytic K_ir_4.1 channels also lead to recovery of basal [K^+^]_o_ and neuronal excitability, particularly during repetitive stimulation, and thereby prevent the emergence of epileptiform activity. Interestingly, we also found that astroglial K_ir_4.1 channels prominently regulate neuronal excitability for slow theta rhythmic activity (3 to 10 Hz), which results from probabilistic firing activity induced by sub-firing stimulation coupled to Brownian noise. In all, these data indicate that astroglial K_ir_4.1 channels acutely regulate [K^+^]_o_ and neuronal excitability during specific patterns of neuronal activity.

## Selective contribution of K_ir_4.1 channels and Na/K ATPases to the activity-dependent astroglial potassium uptake

The model can also be used to investigate the relative contribution of the different current components mediating the activity-dependent astroglial total K^+^ clearance. We thus here comparatively assessed the astrocyte net K^+^ flux mediated respectively by K_ir_4.1 channels and Na/K ATPase during the different regimes of activity (Fig. 4).

**Figure 4.**
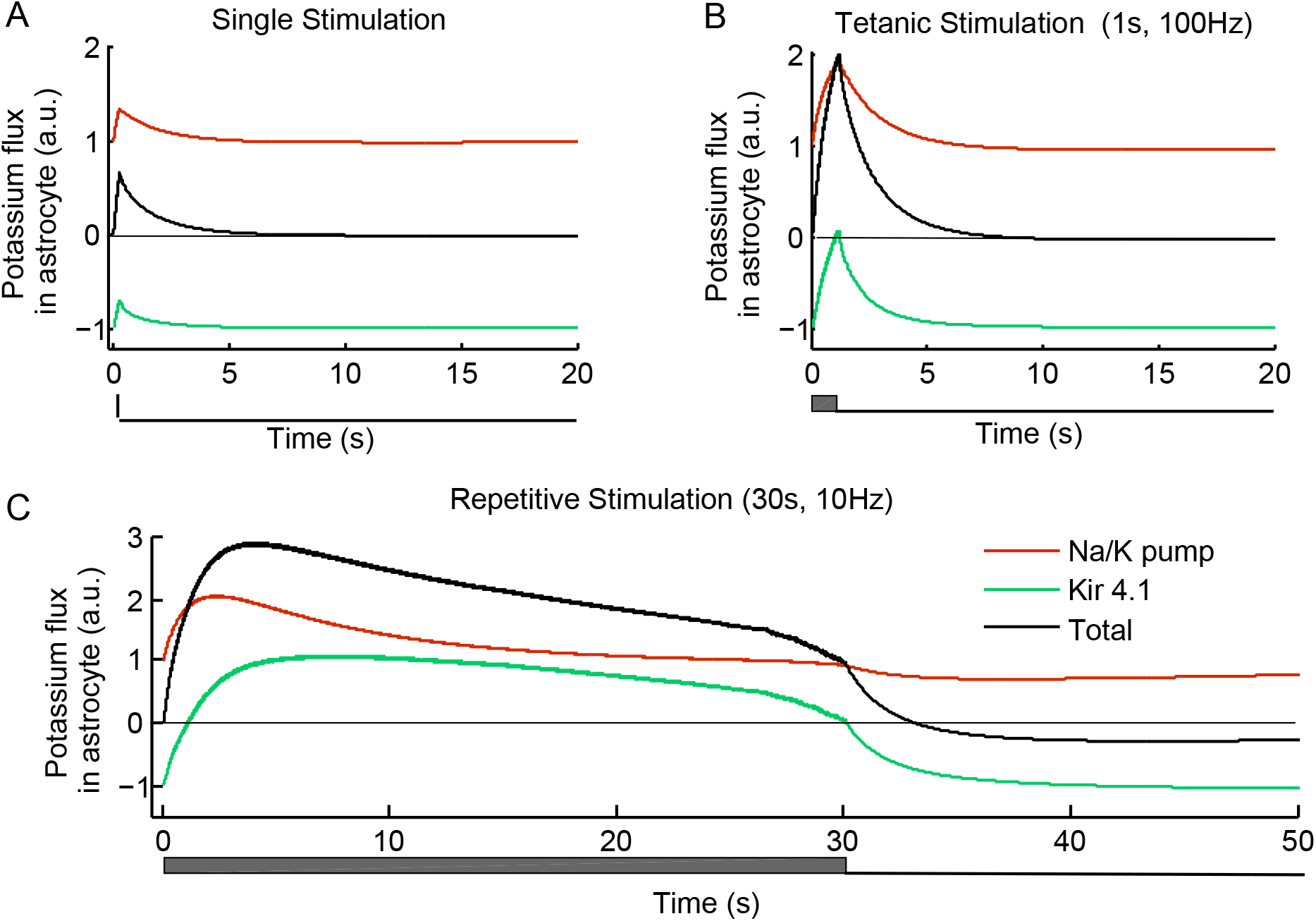
Decomposition of evoked potassium flux in astrocytes. *A)* Plot of astrocyte total K^+^ flux (black), and specific contribution of astroglial K_ir_4.1 channels (green) and Na/K ATPases (red) to the total flux during single afferent stimulation. At rest, the K_ir_4.1 current is outward and counterbalances the inward flux from the Na/K pump. SC stimulation increases both currents, but not sufficiently to reverse the K_ir_4.1 fluxes. *B)* Plot of the same astrocytic K^+^ fluxes during tetanic stimulation (1s, 100Hz). For this level of stimulation, the flux mediated by K_ir_4.1 channels gets outward at the end of the stimulation. *C)* Plot of the same astrocytic fluxes during repetitive stimulation (30s, 10Hz). Here, the K_ir_4.1 fluxes increase and get quickly outward until the end of the repetitive stimulation.

At rest, the K_ir_ current is outward and counterbalances the inward flux mediated by the Na/K ATPase (Fig. 1D). A single afference stimulation increases the activity of Na/K ATPase (Fig. 4A, red) and concomitantly decreases the outward K^+^ flux mediated by the Kir channels, but not sufficiently to reverse it (Fig. 4A green). This combined effect nevertheless significantly and synergistically increases the total K^+^ flux in astrocytes (0.6 a.u. vs 0.3 for Na/K ATPase alone, Fig. 4A, black).

Similar effects on both astrocytic Na/K ATPase and K_ir_4.1 channels-mediated K^+^ currents are observed during tetanic stimulation (Fig 4B, red and green respectively). However, for this level of activity, the flux mediated by K_ir_4.1 channels gets outward at the end of the stimulation. This elicits a rapid increase in the total K^+^ uptake in astrocytes in response to the enhanced [K^+^]_o_ mediated by local neuronal activity (Fig 3A). Thus for this regime of activity, K_ir_4.1 channels and Na/K ATPase similarly contribute to the total astrocytic K^+^ influx.

During repetitive stimulation, the model predicts a two-fold increase in the Na/K ATPase activity (Fig. 4C red), while the K^+^ flux mediated by the K_ir_4.1 channels is significantly stronger than for the other regimes of activity. Such K_ir_4.1-mediated flux gets quickly inward and remains so until the end of the repetitive stimulation (Fig. 4C, green). Consequently the astrocytic total K^+^ influx peaks as early as 3 seconds after the stimulation onset, which translates into a strong astroglial uptake during the whole period of the repetitive stimulus. Thus, although quantitatively most of the net uptake provided by astrocytes is supported by the Na/K ATPase activity (~ 65 %) for this regime of activity (Fig. 4C), the overall contribution of K_ir_4.1 channels to the total astrocytic K^+^ influx (~ 35 %) provides most of the fast adaptation to the net increase in [K^+^]_o_ (Fig 3A).

## Activity-dependent neuronal Na/K ATPase activity

The model can also be used to study the activity of the neuronal Na/K ATPase during the different regimes of activity (single, tetanic and repetitive stimulations). The kinetics of their activity correlates with the slow recovery of neuronal intracellular K^+^ levels [K^+^]_i_ (Fig. 5). It can indeed take 10 seconds for neurons to recover their basal [K^+^]_i_ (Fig. 5). This is due to the long-lasting changes in extracellular and intra-astroglial [K^+^] following neuronal stimulation that have been reported by most studies (Dallerac et al., 2013). Neuronal K^+^ channels and pumps may thus remain activated as long as extracellular K^+^ did not recover to basal levels. However in the model, the stimulations used in our study induce relatively small to moderate changes in neuronal [K^+^]_i_, compared to the high basal neuronal [K^+^]_i_ (135 mM): − 0.4 mM for single stimulation, −1.5 mM for tetanic stimulation, and − 24 mM for repetitive stimulation. Such variations in neuronal [K^+^]_i_ by themselves induce only little changes in the Nernst equilibrium potential for K^+^ (E_K_: −103.7 mV without stimulation, −103.6 mV for single stimulation, −103.3 mV for tetanic stimulation and −98.6 mV for repetitive stimulation).

**Figure 5.**
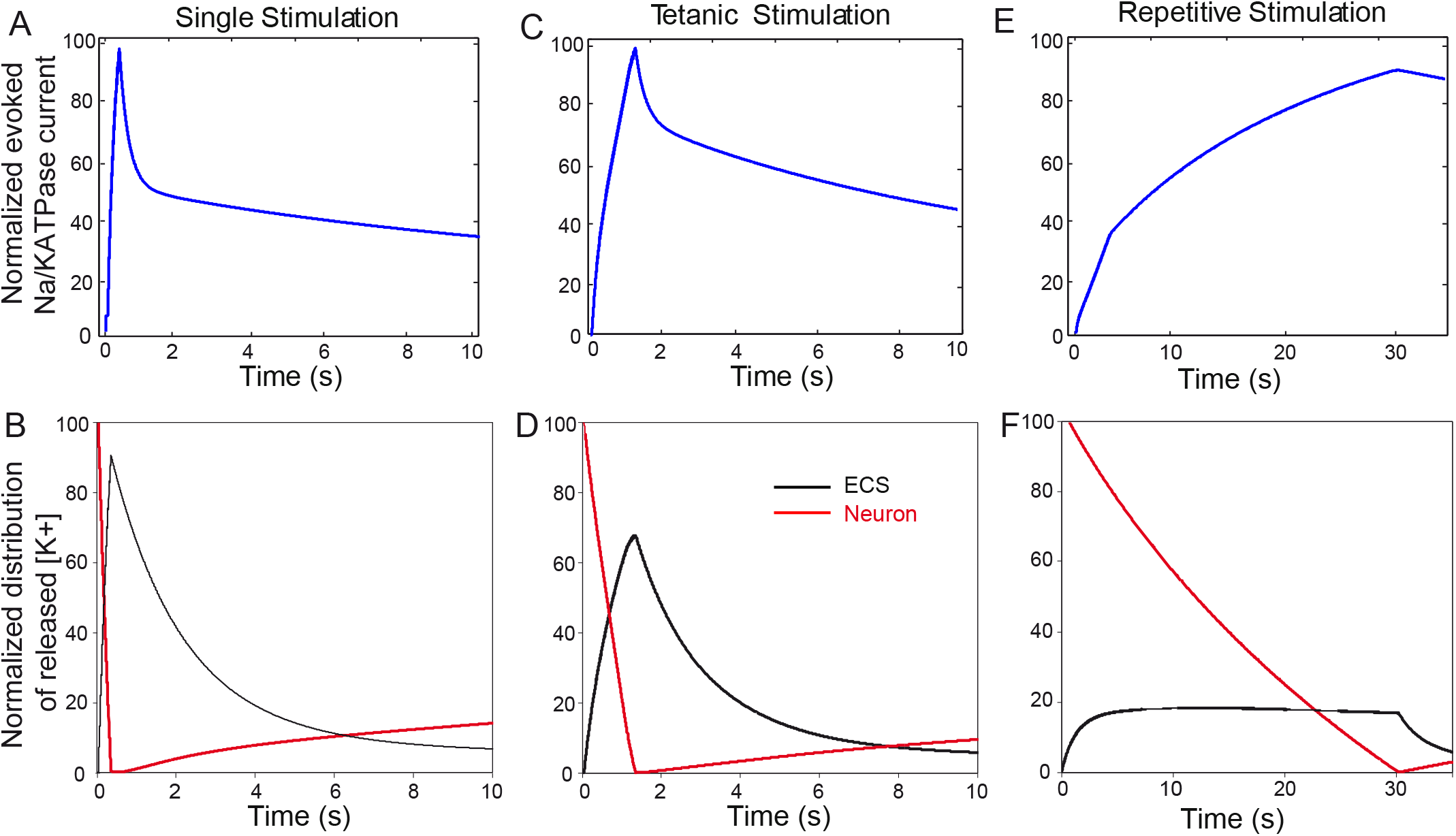
Evoked neuronal Na/K ATPase’s current during the different regimes of activity. *A,C,E)* Normalized neuronal Na/K ATPase current induced by single *(A)*, tetanic (100 Hz, 1 s − *C)* and repetitive (10 Hz, 30 s − E) stimulations. Normalization is performed to Na/K’s current peak amplitude in each regime of activity. *B,D,F)* Normalized K^+^ redistribution between the neuronal and extracellular space compartments induced by single *(B)*, tetanic (100 Hz, 1 s) *(D)* and repetitive (10 Hz, 30 s) *(F)* stimulations. For all regimes of activity, neuronal K^+^ (red) is released in extracellular space (black) during the stimulation initiated at time t=0, and is then cleared by the astrocyte before being redistributed back to the neurons. The redistribution is normalized to the total amount of released K^+^ during each regime of activity.

## Impact of the extracellular space volume on extracellular potassium, astroglial membrane potential and potassium uptake in response to tetanic stimulation

The volumes of the extracellular space, neurons and astrocytes have been reported to represent 20%, 40% and 40 % of the total tissue volume respectively, as assessed by different techniques (Dietzel et al., 1989; Chen and Nicholson, 2000; Magzoub et al., 2009). The model simulations do depend on the values of these volumes. We here investigated the dynamics of [K^+^]_o_ and astroglial membrane potential (Vm) in response to tetanic stimulation (100 Hz, 1s) as a function of different compartment volume proportions (Extracellular space (ECS):neuron:astrocytes) corresponding to the normal situation (20%:40%:40%, Ct ECS, blue), low ECS with cell swelling (11%:44.5%:44.5%, green) and high ECS with cell shrinking (33%:33%:33%, red). The temporal representation of [K^+^]_o_ and astroglial Vm dynamics induced by tetanic stimulation shows that the activity-dependent evoked changes in both parameters are inversely correlated to the size of the ECS (Fig. 6A). The phase diagram illustrating astrocytic K^+^ uptake as a function of activity-dependent changes in [K^+^]_o_ evoked by tetanic stimulation further confirms such result (Fig. 6B).

**Figure 6.**
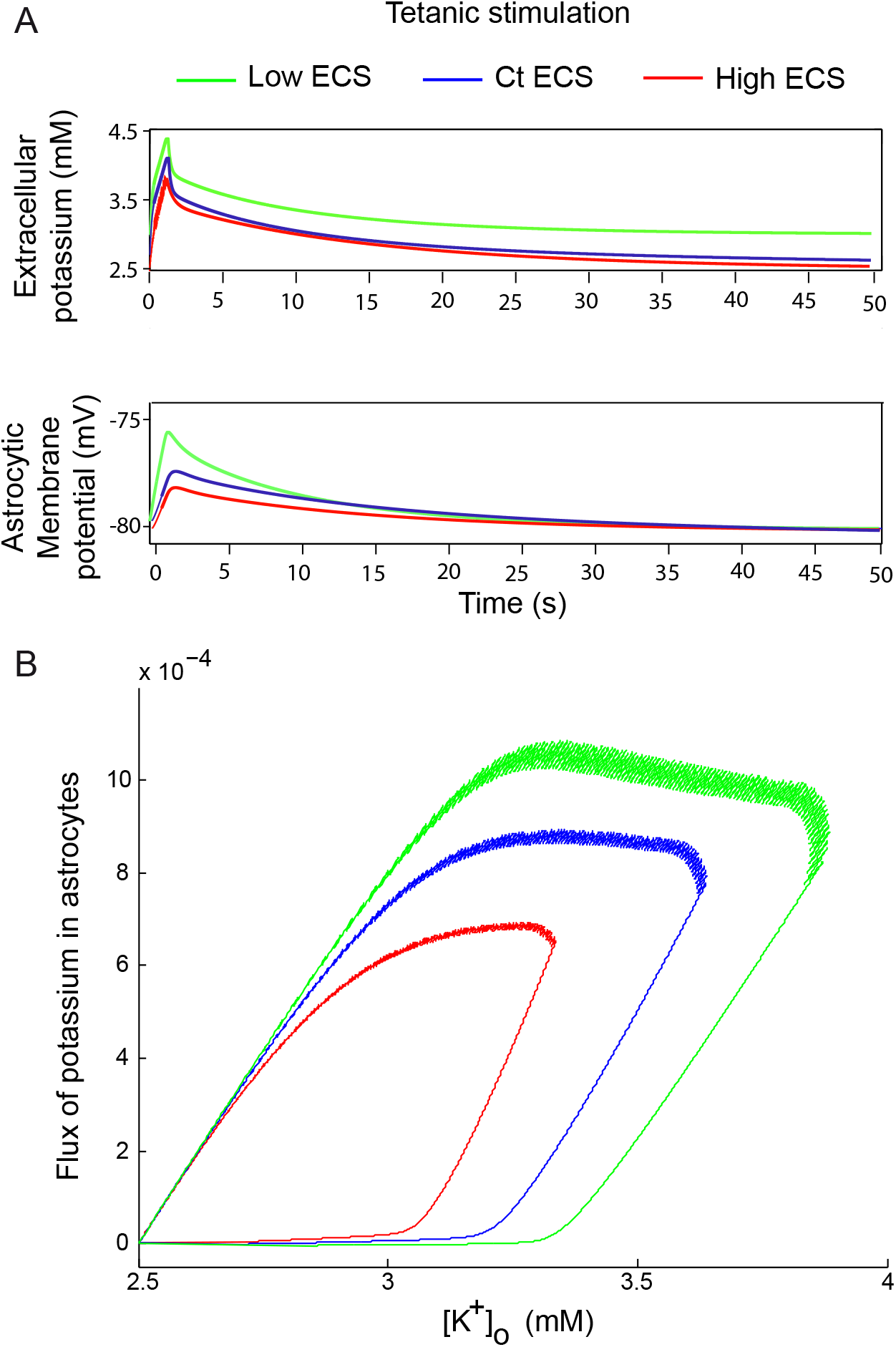
Extracellular potassium, astroglial membrane potential and potassium uptake in response to tetanic stimulation (100 Hz, 1s) as a function of extracellular space volume. *A)* Time-dependent representation of extracellular K^+^ levels and astroglial membrane potential for different compartment volume proportions (ECS / neuron / astrocytes): control situation (20 % / 40 % / 40 % Ct ECS, blue), low ECS (11% / 44.5% / 44.5%, green), high ECS (33% / 33% / 33%, red). *B)* Phase diagram for several extracellular compartment size (Ct, low and high ECS) illustrating astrocytic K^+^ uptake as a function of activity-dependent changes in [K^+^]_o_ evoked by tetanic stimulation.

## Discussion

### Contribution of K_ir_4.1 channel dysfunction to brain diseases: experimental data

Initially, pharmacological studies using non selective blockers of K_ir_4.1 channels, and more recently molecular investigations using glial conditional K_ir_4.1 knockout mice have both suggested an important role for K_ir_4.1 channels in the astrocytic control of [K^+^]_o_ [20–23]. However due to several limitations of these models, mostly related to their lack of molecular (K_ir_4.1), cellular (astrocyte) or temporal selectivity, both approaches do not provide clear information about the specific acute role of astrocyte K_ir_4.1 channels in physiological conditions. They however suggested that disruption of K_ir_4.1 function leads to several defects at the cellular and network levels and can contribute to the development of some pathologies. Indeed, pharmacological blockade of astrocytic K^+^ uptake has reportedly been found to increase neuronal excitability and generates epileptiform activity (Janigro et al., 1997). Remarkably knocking-out K_ir_4.1 channels in a non-conditional manner resulted in a lethal phenotype at postnatal day 20, which was presumably related to myelin development disorders (Neusch et al., 2001) and alteration of ventral respiratory rhythms (Neusch et al., 2006). However, the glial conditional K_ir_4.1 knockout mice also display seizures and die three weeks after birth (Djukic et al., 2007), indicating that alterations in glial K_ir_4.1 have a major impact. At the cellular level, K_ir_4.1^−/−^ astrocytes displayed higher membrane resistance (5 to 20 MΩ) (Djukic et al., 2007), and a depolarized resting membrane potential (~ −60 mV *in vivo* (Chever et al., 2010), ~ −30 mV in brain slices (Djukic et al., 2007)), which likely mediates the loss of synaptically-evoked depolarization in these cells (Djukic et al., 2007). The membrane potential depolarization is likely attributable to the impairment of the K^+^ gradient through the astrocytic membrane. Such hypothesis is however challenged by whole-cell patch clamp experiments, where theoretically both [K^+^]_i_ and [K^+^]_o_ can be controlled. According to the Goldman-Hodgkin-Katz equation, one would indeed expect to be able to restore normal astrocytic membrane potential by manipulating [K^+^]_i_ and [K^+^]_o_ in such mice. The failure in doing so may indicate chronic alterations of other astroglial K^+^ conductances in K_ir_4.1^−/−^ mice. Expectedly, synaptic function is altered in these mice, as shown by decreased spontaneous excitatory postsynaptic currents (Djukic et al., 2007) and increased short-term plasticities, such as post-tetanic potentiation and responses to repetitive stimulation (Sibille et al., 2014). In addition, the decay time of [K^+^]_o_ transients induced by high neuronal activity increases, as shown *ex vivo* in brain slices (Bay and Butt, 2012) or *in vivo* in the cortex of K_ir_4.1^−/−^ mice (Chever et al., 2010). In sum, these initial studies indicate that the K_ir_4.1 channel is probably not the sole, but is definitely the main actor involved in [K^+^]_o_ regulation.

Since then, several studies performed in other models and using different approaches further indicate that the expression of K_ir_4.1 channels and the associated extracellular K^+^ homeostasis are altered in several pathologies. This has been shown for epilepsy (Xiong and Stringer, 1999), ischemia (Pivonkova et al., 2010) or Rett syndrome, a X-linked neurodevelopmental disorder resulting from mutations in methyl-CpG-binding protein 2 (MeCP2) (Kahanovitch et al., 2018). Antibodies against K_ir_4.1 channels have also been suggested as biomarkers for multiple sclerosis and possibly as a cause of the disease via an auto-immune mechanism (Srivastava et al., 2012), although controversies have recently emerged (Filippi et al., 2014; Nerrant et al., 2014; Gu, 2016; Navas-Madronal et al., 2017). Altogether, these data describe mostly correlations between alterations in K_ir_4.1 channel expression and the diseases, but no causal link. Since the physiopathological consequences of the bidirectional communication between neurons and astrocytes are not fully understood, it is indeed still unclear whether alterations in K_ir_4.1 channels and extracellular K^+^ homeostasis cause, worsen, fight these diseases, or rather represent an accompanying phenomenon. Yet a few recent studies have shown that dysfunction of [K^+^]_o_ regulation by astroglial Kir4.1 channels contributes to brain pathologies. Mutations of the human K_ir_4.1 encoding gene KCNJ10 are responsible for the neurodevelopment disorder termed EAST/SESAME syndrom, which includes inherited epilepsy and ataxia (Bockenhauer et al., 2009; Scholl et al., 2009). Deficits in K_ir_4.1 channels functional expression also contributes to dysfunction of striatal medium spiny neurons, including hyperexciblity, and motor coordination in a mouse model of Huntington’s disease (Tong et al., 2014). Finally, K_ir_4.1 channel dysfunction is also involved in psychiatric diseases, since alteration in K_ir_4.1 expression in the lateral habenula drives enhanced bursting activity of neurons and depression-like symptoms in a rat model of depression (Cui et al., 2018). Thus astroglial K_ir_4.1 channels may well represent an alternative therapeutic target for several diseases.

### Computational modeling of astroglial regulation of extracellular potassium levels

Given the high sensitivity of neuronal excitability and synaptic activity to [K^+^]_o_, getting insights into the astroglial contribution to the dynamic redistribution of ionic fluxes during neurotransmission in physiological conditions is of major interest. K^+^ currents evoked by synaptic activity have unanimously been characterized as the primary astrocyte current in various brain areas (Dallerac et al., 2013). In addition, the direct correlation between activity-dependent increase in [K^+^]_o_ and astrocytic K^+^ entry has been proved using synchronous ion sensitive electrodes and astrocytic membrane potential recordings in brain slices (Neprasova et al., 2007) and *in vivo*, in particular during epileptiform activity or specific brain rhythms (Amzica and Steriade, 2000). *In vivo* data also report that the timing of extracellular K^+^ clearance is strongly correlated with the latency of the astrocytic depolarization (Amzica and Steriade, 2000; Amzica et al., 2002; Chever et al., 2010). Thus in response to stimulation, neurons display a fast transient activity inducing a delayed depolarization of astrocytic membrane potential concomitant to increases in [K^+^]_o_ (Amzica and Steriade, 2000; Chever et al., 2010). Similarly, in hippocampal acute slices, SC stimulation induces an activity-dependent potentiation of [K^+^]_o_, which is strongly correlated to similar potentiation of astrocytic membrane depolarizations, here assessed by evoked currents (Meeks and Mennerick, 2007).

However, as previously mentioned, the currently available experimental pharmacological and molecular approaches do not permit to assess quantitatively the specific and acute role of individual astroglial channels such as K_ir_4.1 in the activity-dependent redistribution of K^+^ fluxes within the neuroglial and extracellular compartments. Computational modeling is an alternative approach of choice to address this issue that we and others have developped.

In the early 80s, Gardner-Medwin was among the first to study the properties of extracellular K^+^ diffusion using numerical simulations. He showed that K^+^ diffusion in the ECS alone does not account for the experimental recordings of sink and source K^+^ fluxes in cortical areas following brain stimulation (Gardner-Medwin, 1983). He indeed calculated that K^+^ diffusion in the ECS accounts for only ~ 25% of the observed K^+^ flow throughout cortical layers. This led him to postulate the existence of “transfer cells” that would be responsible for the remaining 75% of the measured ionic fluxes (Gardner-Medwin, 1983; Dietzel et al., 1989). This has initially been confirmed in the retina by modeling extracellular K^+^ diffusion, which was shown to be more efficient in the presence of Müller cells (Newman et al., 1984). Since then, astrocytes have experimentally been identified as these “transfer cells”, notably as they express high levels of K_ir_4.1 channels and Na/K ATPases. As described above, a number of computational models have investigated the astroglial regulation of [K^+^]_o_ and their impact on neuronal activity. However most of them focused on pathological conditions such as epilepsy (Cressman et al., 2009; Ullah et al., 2009). In the latter studies, a tri-compartment (neuron, astrocyte and extracellular space) is included and the astrocytic K^+^ dynamics is modeled by a unidirectional flux from the ECS into the astrocytes. Thus the astrocytic effect is simply modeled by a generic astroglial uptake of K^+^, and independently of the molecular mediators. However, other models did incorporate bidirectional K^+^ flux within astrocytes, in particular during pathological situations such as spreading depression (Kager et al., 2000). Such type of model, which also includes three compartments but does not take into account changes in astrocyte membrane potential, does capture a long-lasting dynamic increase in [K^+^]_o_, following brief neuronal activations. Other models performed in a pathological context however included more complexity and did take into account not only specific ion channels and transporters, but also variations in astrocyte membrane potential and volume induced by strong ionic variations occurring during pathological situation such as stroke (Dronne et al., 2006; Dronne et al., 2007). Such approach showed that astrocytes can control extracellular levels of K^+^ and glutamate solely during mild ischemia. Interestingly, more recent modeling studies on astrocyte ion dynamics and implications of spatial K^+^ buffering put forward a multiscale theory that includes not only local ionic exchanges and diffusive processes, but also osmotic changes and volume regulations (Ostby et al., 2009; Oyehaug et al., 2012; Halnes et al., 2013; Halnes et al., 2016). This model indeed proposes that astroglial ion dynamics might modulate microscopic liquid flow via osmotic effects resulting in whole-brain macroscopic flow. In this physical model, the local ionic changes occurring during neurotransmission are described by the Kirchhoff-Nernst-Planck equations, which account for electrodiffusion in the limited compartments where ions are actually released. In addition, ionic diffusion between astrocytes is mediated by gap junction channels, which create narrow passages, leading to a Poissonian law for the exchange rate (Holcman and Schuss, 2013). Such local description emphasizes the global electrodiffusion here simulated during either dynamical changes induced by neuronal activity or during subsequent steady states (Ostby et al., 2009; Oyehaug et al., 2012; Halnes et al., 2013; Halnes et al., 2016). Osmotic pressures and aquaporin4 (AQ4) water based fluxes are also simulated and are proposed to account for changes in both astrocytic swelling and ECS volume. Altogether, this study elegantly proposes a mathematical description of the ionically-driven fluxes of water and ions described as “Potassium Buffering and Liquid flow”. The model is nevertheless limited by the facts that the different key elements taken into account are simulated independently, and that astrocytes are approximated by cylinders, neglecting the spatially round astrocytic structure that deviate significantly from cable.

In all, modeling of ion neuroglial interactions has primarily been focused on regulation of [K^+^]_o_ in pathological conditions. Over the years models have tremendously evolved to incorporate multiscale key parameters to account for the endogenous complexity of cellular interactions in micro-compartments.

### Which role for astrocytes in the dynamics of ion redistribution during physiological neurotransmission?

However, modeling studies did not examine the synchronous activities of neurons and astrocytes together with [K^+^]_o_ variations during physiological neurotransmission, while taking into account astrocyte bidirectional K^+^ fluxes, the contribution of specific astroglial channels and the dynamics of their membrane potential during evoked activity. We have thus developed a modeling approach (Sibille et al., 2015) to achieve such a goal, and have initially focused on the role of astroglial K_ir_4.1 channels in the neuroglial K^+^ cycle during neurotransmission, as summarized in the present article. We thereby found that astrocyte K_ir_4.1 channels play a major role in clearance of extracellular K^+^, dynamics of astrocyte and neuron membrane potential, especially during high neuronal activity, and robustly modulate neuronal excitability for slow rhythmic activity.

In contrast to action potentials, which are characterized by a fast dynamics of a few milliseconds, astrocyte K^+^ buffering lasts tens of seconds. Our model showed that the factors controlling the slow timescale of astroglial K^+^ clearance are K_ir_4.1 channels. Our model also confirms that the major effect of acute K_ir_4.1 channels disruption consists in a slower extracellular K^+^ dynamics, as experimentally recorded *ex vivo* and *in vivo* in several brain regions (Chever et al., 2010; Haj-Yasein et al., 2011; Bay and Butt, 2012). However, we also found that K_ir_4.1 channels alter [K^+^]_o_ peak amplitudes during repetitive stimulations, suggesting that K_ir_4.1^−/−^ mice might have developed compensatory mechanisms to maintain homeostasis of [K^+^]_o_. Moreover, the redistribution of synaptically-released K^+^ during different regimes of activity indicates that the higher the activity, the lower the proportion of released K^+^ remains transiently in the extracellular space. This indicates that K_ir_4.1 channels have a strong clearance capacity, in particular during high regimes of activity, where [K^+^]_o_ can reach up to ~ 5 mM. We also found that astrocytic K_ir_4.1 channels are sufficient to account for the overall astrocytic membrane potential dynamics during neuronal activity, confirming the experimental data obtained in astrocytes from the K_ir_4.1^−/−^ mice (Djukic et al., 2007).

Finally, our model reveals that K_ir_4.1 channels strongly modulate action potential discharges specifically during certain regimes of activity, such as repetitive stimulation. K_ir_4.1channels indeed play a major role in the regulation of [K^+^]_o_ during this regime of activity, most likely because such stimulation results in a sustained, but moderate increase in [K^+^]_o_ of ~ 6 mM for ~ 20 seconds compared to the capacity of astrocytes to buffer up to ~ 14 mM of [K^+^]_o_ during this regime. The specificity of such astroglial regulation of neuronal firing is further illustrated by the fact that astroglial K_ir_4.1 channels have almost no impact on firing induced by single and tetanic stimulations. This most likely stems from the only transient and small increase in [K^+^]_o_ reaching ~ 2.5 mM 450 milliseconds after the single stimulation, and ~ 3.5 mM 1.5 second after the tetanic stimulation. Nevertheless we found a prominent and specific contribution of K_ir_4.1 channels in the probabilistic firing activity induced by 3 to 10 Hz sub-firing stimulations, suggesting a key role of these channels during sustained theta rhythmic activity, the hallmark of *in vivo* active behavior. Noteworthy, these data imply that K_ir_4.1 channels can finely tune action potential discharges involving low, but long-lasting, increase in [K^+^]_o_.

Noteworthy, our model revealed that specific and acute inhibition of K_ir_4.1 channels slows down, but does not abolish, astroglial uptake of excess K^+^ during single, tetanic and repetitive stimulations, suggesting the contribution of additional channels or pumps in astrocytes. We thus here used our model to generate new predictions, and in particular to investigate the relative contribution and dynamics of K_ir_4.1 channels and Na/K ATPases, the two types of channel/pump included in our model, to the astroglial total K^+^ clearance during the different regimes of activity. We found that Na/K ATPases contribute significantly to K^+^ clearance, as reported experimentally (Larsen et al., 2014), and that their contribution is activity-dependent. Indeed, while K_ir_4.1 channels and Na/K ATPases contribute equally to the total astrocyte K^+^ influx during single and tetanic stimulation, most of the uptake (~65%) is provided by Na/K ATPases during repetitive stimulation. However, the relative activity-dependent variation in K^+^ flux mediated by K_ir_4.1 channels is stronger than the one of Na/K ATPase for this regime activity. This indicates that the sensitivity of K_ir_4.1 channels to activity, reflected by increase in [K^+^]_o_, is higher than the one of Na/K ATPases.

As an element of comparison, we here also analyzed the Na/K ATPase activity of neurons during the different regimes of activity, as well as the total K^+^ uptake capacity of neurons following release. The kinetics of Na/K ATPase activity is slow and correlates with the slow recovery of neuronal [K^+^]_i_. It can indeed take several seconds for neurons to restore their basal [K^+^]i, most likely due to the prolonged increase in [K^+^]_o_ following neuronal activity, which maintains prolonged activity of Na/K ATPases.
In fact, the activity-dependence of the neuronal Na/K pump ATPase is limited: as found in the retina (Reichenbach et al., 1992), the neuronal Na/K ATPase activity is not changing with [K^+^]_o_ increase above 3mM. However, adding slower timescale K^+^-dependent conductances in our neuron model should induce faster recovery of neuronal [K^+^]_i_ and may be of interest to implement in future development of the model. This poor ability of neurons to uptake the released K^+^ is actually found for all regimes of activity. Indeed 10 seconds after the tetanic stimulation, the neuron has only taken up 10 % of the released K^+^. By this time, the astrocyte has already buffered up to 90 % of the released K^+^, so that [K^+^]_o_ is back to basal levels. During repetitive stimulation, only 20 % of the activity-induced K^+^ release remains in the extracellular space, indicating that the remaining 80 % have been taken up directly by the astrocyte. The neuronal ability to uptake synaptically-released K^+^ is thus a limiting factor slowing down the neuroglial K^+^ cycle during neurotransmission.

Finally, we found that the sizes of the extracellular space and of the cellular compartments are critical parameters which strongly influence the dynamics of [K^+^]_o_ induced by neuronal activity. The activity-dependent changes in [K^+^]_o_ are indeed inversely correlated to the size of the ECS. This suggests that changes in neuronal activity or pathological conditions, both known to alter cell size and consequently ECS, can by themselves impact the dynamics of neuroglial K^+^ cycle. Because the size of the ECS is much smaller than the one of the cellular compartments (20 % / 80 %, respectively), accumulation of a small number of released ions in a reduced ECS should still significantly increase the local ionic concentrations. Such effect should not only depend on the quantity of released ions, but also on the diffusion and tortuosity properties of the ECS. Considering such elements in future development of our model is of interest. Yet the swelling of astroglial processes following neuronal activity and known to reduce ECS volume, remains poorly investigated, most likely due to the limited tools to assess such phenomenon in living tissues. The high co-localization of K_ir_4.1 and AQ4 channels within astroglial processes however suggests co-regulation of their distribution and functionality. In fact both channels influence each other, since AQ4 deletion impairs evoked [K^+^]_o_ (Strohschein et al., 2011), while genetic or pharmacological invalidation of K_ir_4.1 channels blocks the activity-induced swelling in the spinal cord (Dibaj et al., 2007). Interestingly, it has been hypothesized that AQ4-dependent volume regulation could be coupled to K^+^ regulation (Amiry-Moghaddam et al., 2003). This hypothesis is however subject to controversies, since AQ4 deletion facilitates K^+^ buffering in the hippocampus (Strohschein et al., 2011), but K_ir_4.1 deletion does not impairs neurovascular coupling (Metea et al., 2007). Altogether, the possibility that membrane potential-dependent K^+^ fluxes may regulate osmotic pressure changes is an interesting hypothesis, but a direct link between these phenomena still remains largely elusive.

### Perspectives

In summary, we have here described a simplified tri-compartment model accounting for K^+^ redistribution between neurons, astrocytes and the extracellular space during neurotransmission. Such model includes the minimal set of channels and pumps in both neurons and astrocytes, which accounts for experimental data related to activity-dependent changes in [K^+^]_o_ and cell membrane potential. However this tri-compartment model, as most current models, did not account for the sophisticated multiscale geometry of astrocytes and neurons, as well as for the diversity of ionic channels present on their membranes. Calcium-dependent K^+^ channels (K_Ca_3.1), but also other channels, transporters or exchangers (such as Cx hemichannels, Na^+^/K^+^/Cl^−^ co-transporter (NKCC1) K^+^/C^−^ exchanger, glutamate transporters) (Kindler et al., 2000; Dronne et al., 2006; Verkhrastky and Butt, 2013) could indeed also play a role in the regulation of activity-dependent changes in [K^+^]_i_ or [K^+^]_o_. Thus incorporating in the future additional channels as well as the complex geometries of cellular nanodomains will be of particular interest.

Furthermore, several questions remain: 1) How localized is the regulation of K^+^ dynamics: is the K^+^ regulated locally along dendrites, axons or at synapses? 2) How astrocytes spatially redistribute K^+^ ions released by neurons? 3) How to generalize the model discussed here to integrate the complex geometry of neurons and astrocytes? 4) What is the specific contribution of other ionic neuronal and astroglial channels? 5) Can the astroglial regulation of [K^+^]_o_ in physiological conditions alter the presynaptic vesicular release? 6) Does and how the extracellular K^+^ dynamics control astroglial calcium signaling? Future experimental and modeling studies should provide further insights into these issues.

## Acknowledgments

This work was supported by grants from European Research Council (Consolidator grant N°683154) and Lejeune Foundation to N.R., and the doctoral school “Frontiers in Life Science”, Paris Diderot University, Bettencourt Schuller foundation to J.S..

